# Distinct microbial communities along the chronic oil pollution continuum of the Persian Gulf converge with oil spill accidents

**DOI:** 10.1101/2020.07.25.221044

**Authors:** Maryam Rezaei Somee, Seyed Mohammad Mehdi Dastgheib, Mahmoud Shavandi, Leila Ghanbari Maman, Kaveh Kavousi, Mohammad Ali Amoozegar, Maliheh Mehrshad

## Abstract

Persian Gulf hosting *ca.* 48% of the world’s oil reserves; has been chronically exposed to natural oil seepage. Oil spill events have been studied over the last decade; however, the influence of chronic oil exposure on the microbial community of the Persian Gulf has remained unknown. We performed genome-resolved comparative analyses of the water and sediment’s prokaryotic community along the Gulf’s pollution continuum (Strait of Hormuz, Asalouyeh and Khark Island). The continuous exposure to trace amounts of pollution has shifted the microbial profile toward the dominance of *Oceanospirillales, Flavobacteriales, Alteromonadales*, and *Rhodobacterales* in Asalouyeh and Khark samples. Intrinsic oil-degrading microbes present in low abundances in marine habitats; experience a bloom in response to oil pollution. Comparative analysis of the Persian Gulf samples with 106 oil-polluted marine samples reveals the pollutant’s hydrocarbon content, exposure time and sediment depth as main determinants of microbial response to pollution. High aliphatic content enriches for *Oceanospirillales, Alteromonadales* and *Pseudomonadales* whereas, *Alteromonadales, Cellvibrionales, Flavobacteriales* and *Rhodobacterales* dominate polyaromatic polluted samples. In sediment samples, *Deltaproteobacteria* and *Gammaproteobacteria* had the highest abundance. In chronic exposure and oil spill events, the community composition converges towards higher dominance of oil-degrading constituents while promoting the division of labor for successful bioremediation.

**Originality-Significance Statement:** The impact of anthropogenic oil pollution on the microbial community has been studied for oil spill events; while the influence of long-term chronic exposure to oil derivatives on The microbes has remained unknown. Persian Gulf hosts ca. 48% of the world’s oil reserves and has been chronically exposed to natural and accidental oil pollutions. Different pollutant profilesin different locations and the recurrent pollution events; make Persian Gulf an ideal model system to analyse the impact of oil hydrocarbon on the microbial community and the recovery potential of marine ecosystems after pollution. In this study we perform an extensive analysis of thhe Persian Gulf’s water and sediment samples along the water circulation and pollution continuum for the first time. Our results show that these long-standing trace exposure to oil has imposed a consistent selection pressure on the Gulf’s microbes; developing unique and distinct communities along the pollution continuum. Our extensive genome-resolved analysis of the metabolic capabilities of the reconstructed MAGs shows an intricate division of labor among different microbes for oil degradation and determine the major drivers of each degradation step. Intrinsic oil-degrading microbes (e.g., *Immundisolibacter, Roseovarius* and *Lutimaribacter*) bloom along the Persian Gulf’s pollution continuum and function as the main oil degraders. Comparative study of PG datasets with 106 oil-polluted marine samples (water and sediment) reveals similar community compositions in the Persian Gulf’s water and sediment samples to those of oil spill events and suggests hydrocarbon type and exposure time as the main determinants of the microbial response to oil pollution.

## Introduction

Exposure to oil and gas derivatives in marine ecosystems rich in oil reservoirs is inevitable due to natural seepage. Intensive industrial oil exploration and transit over the last century have further increased the risk of pollution in these ecosystems (Brussaard *et al.*, 2016). Persian Gulf is a relatively shallow evaporative basin (Hajrasouliha and Hassanzadeh, 2015) that hosts more than 48% of the world’s oil reservoirs (Joydas *et al.*, 2017). The largest recorded oil spill in the Persian gulf dates back to 1991 (Chatterjee, 2015). However, this ecosystem has been chronically exposed to oil hydrocarbon pollution through natural seepage, accidental oil derivatives release from transit tankers or refinery facilities, and discharge of oily wastes and heavy metals from offshore drilling sites. Additionally, the limited water circulation in this semi-enclosed habitat (Ludt *et al.*, 2018) prolongs the residence time of pollutants in the basin. Since oil is of biogenic origin, a wide variety of microbes are capable of its degradation (Delille *et al.*, 2002). While representing low abundances in the pristine marine community, these intrinsic oil-degrading taxa bloom in response to oil pollution and play a critical role in the bioremediation process (Yergeau *et al.*, 2015). Because of their vigilance in responding to pollution, they could be considered as microbial indicators of trace oil pollution (Hu *et al.*, 2017).

In the case of oil seepage or spill accidents, the natural marine microbial community exposed to oil hydrocarbons; responds with fluctuating composition as different oil components are gradually degraded (Ludt *et al.*, 2018). In the Persian Gulf, as the input water is carried along the Gulf’s water circulation, it is exposed to different types of oil derivatives in different locations. Sporadic cultivation efforts have isolated Naphthalene degrading (*Shewanella, Salegentibacter, Halomonas, Marinobacter, Oceanicola, Idiomarina* and *Thalassospira*) (Hassanshahian and Boroujeni, 2016) and crude oil utilizing (*Acinetobacter, Halomonas, Alcanivorax, Marinobacter, Microbacterium, Rhodococcus* (Hassanshahian *et al.*, 2012), and *orynebacterium*(Hassanshahian *et al.*, 2014)) bacteria from the Persian Gulf water and sediment samples and a single 16S rRNA amplicon study of the Mangrove forest’s sediment show a community dominated by *Gammaproteobacteria* and *Flavobacteriia* (Ghanbari *et al.*, 2019). However, the impact of chronic exposure to oil derivatives and recurring pollutions on the microbiome of the Persian Gulf, their oil bioremediation capability, and recovery potential remains largely unknown.

To explore the dynamics of the input microbial community along the pollution continuum and their metabolic capability for oil bioremediation; in this study we performed genome-resolved comparative analyses of the Persian Gulf’s water and sediment samples. We collected water and sediment samples from the Gulf’s input water at the Strait of Hormuz. Along the water circulation current, we have also collected samples close to Asalouyeh and Khark Island. Asalouyeh hosts a wide variety of natural gas and petrochemical industries and its surrounding water is exposed to aromatic compounds pollution (Delshab *et al.*, 2016). Khark Island is the most important oil transportation hub of the Persian Gulf continuously receiving major oil pollution (Mirvakili *et al.*, 2013). Our results show a patchy microbial community with contrasting composition and metabolic capabilities along the Persian Gulf’s pollution continuum. Additionally; we compare Persian Gulf’s microbial community with the publicly available metagenomes of oil-polluted marine water (*n*=41) and sediment (*n*=65) samples. Our analyses show that the microbial community of the Persian Gulf is severely impacted by chronic oil pollution and suggest a critical role for hydrocarbon type in defining the bacterial and archaeal community composition of water samples in response to the chronic oil exposure while in sediments the sampling depth is an additional factor.

## Results and discussion

### Persian Gulf water and sediment samples along the oil pollution continuum

Water and sediment samples were collected along the circulation current of the Persian Gulf from Hormuz Island (HW and HS), Asaluyeh area (AW and AS) and Khark Island (KhW and KhS) (**Supplementary Figure S1**). Detailed description of sampling points, their physicochemical characteristics and Ionic content are presented in **Supplementary Tables S1**. No obvious differences in physicochemical characteristics (temperature, pH and salinity) and ionic concentrations were detected among water samples except for the slightly higher salinity of the AW (4%). The GC-FID analyses showed high TPH and PAH concentrations in the Khark sediment sample (KhS) (**Supplementary Table S2**). GC-SimDis analysis showed that >C_40_ hydrocarbons were dominant in the KhS (~14%) followed by C_25_-C_38_ hydrocarbons (**Supplementary Figure S2**). Chrysene, fluoranthene, naphthalene, benzo(a)anthracene and phenanthrene were respectively the most abundant PAHs in KhS. This pollution originates from different sources including oil spillage due to Island airstrikes during the imposed war (1980-1988), sub-sea pipeline failures and discharge of oily wastewater or ballast water of oil tankers (ongoing for ~50 years)(Akhbarizadeh *et al.*, 2016). The concentration of oil derivative pollutants in other water and sediment samples was below our detection limit (<50 μg/L and 1 μg/g respectively). qPCR-mediated estimates of the 16S rRNA gene copy number show an increase in the proportion of bacteria from HW to AW and KhW (87 to 96%) for water samples and from AS to KhS and HS (74 to 99%) for sediment samples obviously, the archaeal communities represent a reverse trend reaching highest proportion in AS (~25%) **(Supplementary Figure S3 & Supplementary Table S1)**.

### Distinct Prokaryotic community composition along the oil pollution continuum of the Persian Gulf

In pristine marine environments, the internal feedback mechanisms of the microbial communities facilitate keeping a steady “average” composition despite changes in factors such as temperature, nutrient supply, and physical mixing (Fuhrman *et al.*, 2015). However, in the Persian Gulf, the continuous exposure to oil pollution in water samples causes spatial patchiness and a shift in the microbial community composition representing a distinct composition along the pollution continuum. The input water (HW) has a typical marine microbial community dominated by *Synechococcales*, SAR11, SAR86, *Flavobacteriales, Actinomarinales* and *Rhodobacterales* (Fuhrman *et al.*, 2015; Salazar and Sunagawa, 2017) whereas along the pollution continuum the community shifts towards a higher relative abundance of *Gammaproteobacteria* and *Bacteroidetes* representatives (**Supplementary Figure S4**). The relative abundance of phyla *Cyanobacteria, Actinobacteria, Marinimicrobia* and order *Cytophagales* of the phylum *Bacteroidetes* decreases, inversely, the relative abundance of phyla *Proteobacteria, Epsilonbacteraeota*, and *Firmicutes* as well as orders *Flavobacteriales* and *Balneolales* of the phylum *Bacteroidetes* consistently increase from HW to AW and KhW (**Supplementary Figures S4**). The order *Synechococcales* negatively responds to oil pollution in marine surface water (Bacosa *et al.*, 2015) and its decrease along the oil pollution continuum from HW to AW and KhW complies with our chlorophyll-a measurements (0.24, 0.091, and 0.013 μg/L respectively) (**Supplementary Table S1**). While oil pollution of the KhW and AW samples was below our detection limit (50 μg/L), their dominant prokaryotic community is remarkably similar to other oil-polluted marine samples (Yergeau *et al.*, 2015). *Oceanospirillales, Flavobacteriales, Alteromonadales* and *Rhodobacterales* have the highest relative abundance in KhW and the prokaryotic community of AW is mainly comprised of *Alteromonadales*, SAR86, *Flavobacteriales, Rhodobacterales* and *Thermoplasmata* (**Supplementary Figure S4**).

In response to oil contamination (e.g. in form of an oil spill) relative abundance of oil-degrading microbes increases (Rodriguez-r *et al.*, 2015) (e.g., *Oceanospirillales, Flavobacteriales, Alteromonadales*, SAR86, and *Rhodobacterales*)(King *et al.*, 2015; Yergeau *et al.*, 2015; Doyle *et al.*, 2018). SAR86 clade is among dominant marine bacteria reported to contain cytochrome P450 and dioxygenase genes that are involved in degrading aliphatic and aromatic xenobiotic compounds (Dupont *et al.*, 2012). SAR86 representatives reach the highest relative abundance in AW where they are exposed to aromatic pollutants. *Alteromonadales* representatives encode enzymes for degrading recalcitrant and toxic branched-chain alkanes, and PAHs thus are mainly involved in the final steps of the degradation process (Das and Chandran, 2011; Hu *et al.*, 2017). They show the highest relative abundance in AW where most of the oil contaminants are low molecular weight aromatic compounds (**Supplementary Figure S4**).

*Oceanospirillales* comprise ca. 20% of the KhW prokaryotic community (compared to ca. 1% in HW and AW). They prevail following marine oil pollution and are involved in the degradation of labile compounds such as non-branched alkanes and cycloalkanes (Mason *et al.*, 2014). *Oceanospirillales* prevalence in KhW suggests a recurring recent pollution.

In sediment samples, the KhS represents a distinct microbial profile from HS and AS. *Alteromonadales, Rhodobacterales, Oceanospirillales, Deferribacterales, Halothiobacillales* and *Balneolales* (>2%) representatives are enriched in this sample (**Supplementary Figure S5**). The main HC pollutants in the Asalouyeh are low molecular weight aromatic compounds that mainly influence the prokaryotic population in the water column and rarely precipitate into sediments hence the similarity of AS to HS microbial composition as they both experience low pollution rates.

Apart from oil-degrading *Proteobacteria* (e.g. *Alteromonadales, Rhodobacterales,* and *Oceanospirillales*), a diversity of sulfur/ammonia-oxidizing chemolithoautotrophic *Proteobacteria* are present in these sediments although at lower abundances e.g., (*Acidithiobacillales* (KhS 1.8%), *Chromatiales* (HS 1.5, AS 1.1, KhS 0.85%), *Ectothiorhodospirales* (HS 3.75, AS 2.3, KhS 1.7%), *Halothiobacillales* (KhS 2.6%), *Thiotrichales* (HS 1.5, AS 1.1, KhS 0.3%), *Thiohalorhabdales* (HS 0.7, AS 1.2, KhS 0.5%), *Thiomicrospirales* (KhS 1.5%)).

Sulfate-reducing bacteria (SRB) in HS comprise up to 16.2% of the community (*Desulfobacterales,* NB1-j, *Myxococcales, Syntrophobacterales* and uncultured *Thermodesulfovibrionia*). Similar groups along with *Desulfarculales* comprise the SRB functional guild of the AS (~18.9%). Whereas *Desulfuromonadales* and *Desulfobacterales* are the SRB representatives in KhS with a total abundance of only ~ 3.3%. The lower phylogenetic diversity and community contribution of SRBs in KhS hint at potential susceptibility of some SRBs to oil pollution or that they might be outcompeted by HC degraders (e.g., *Deferribacterales*). Additionally, KhS is gravel-sized sediment (particles ≥4 millimeters diameter), whereas HS and AS samples are silt and sand-sized sediments (de Sousa *et al.*, 2019). The higher oxygen penetration in gravel particles of KhS hampers anaerobic metabolism of sulfate/nitrate-reducing bacteria hence their lower relative abundance in this sample (**Supplementary Figure S5**).

### Chronic exposure to oil pollution shapes similar prokaryotic communities as oil spill events

We have analyzed the prokaryotic community composition of 41 oil-polluted marine water metagenomes (different depths in the water column) from Norway (Trondheimsfjord), Deepwater Horizon (Gulf of Mexico), the northern part of the Gulf of Mexico (dead zone) and Coal Oil Point of Santa Barbara; together with 65 oil exposed marine sediment metagenomes (beach sand, surface sediments and deep-sea sediments) originating from DWH Sediment (Barataria Bay), Municipal Pensacola Beach (USA) and a hydrothermal vent in Guaymas Basin (Gulf of California) in comparison with the PG water and sediment samples (in total 112 datasets) (**Supplementary Table S3**). This extensive analysis allows us to get a comparative overview of the impact of chronic oil pollution on the prokaryotic community composition.

Hydrocarbonoclastic bacteria affiliated to *Oceanospirillales, Cellvibrionales* (*Porticoccaceae* family) and *Alteromonadales* (Gutierrez, 2017) comprised the major proportion of the prokaryotic communities in samples with higher aliphatic pollution e.g. DWHW.BD3 (sampled six days after the incubation of unpolluted water with Macondo oil), DWHW.he1 and DWHW.he2 (oil-polluted water samples incubated with hexadecane), DWHW.BM1, DWHW.BM2, DWHW.OV1 and DWHW.OV2 (sampled immediately after the oil spill in the Gulf of Mexico) (**Figure 1**). Samples treated with Macondo oil, hexadecane, naphthalene, phenanthrene and those taken immediately after the oil spill in the Gulf of Mexico have significantly lower proportion of SAR11 due to the dominance of bloom formers and potential susceptibility of SAR11 to oil pollutants (**Figure 1**). Our results suggest that samples with similar contaminants and exposure time to oil pollution enrich for similar phylogenetic diversity in their prokaryotic communities (**Figure 1**).

**Figure 1.**
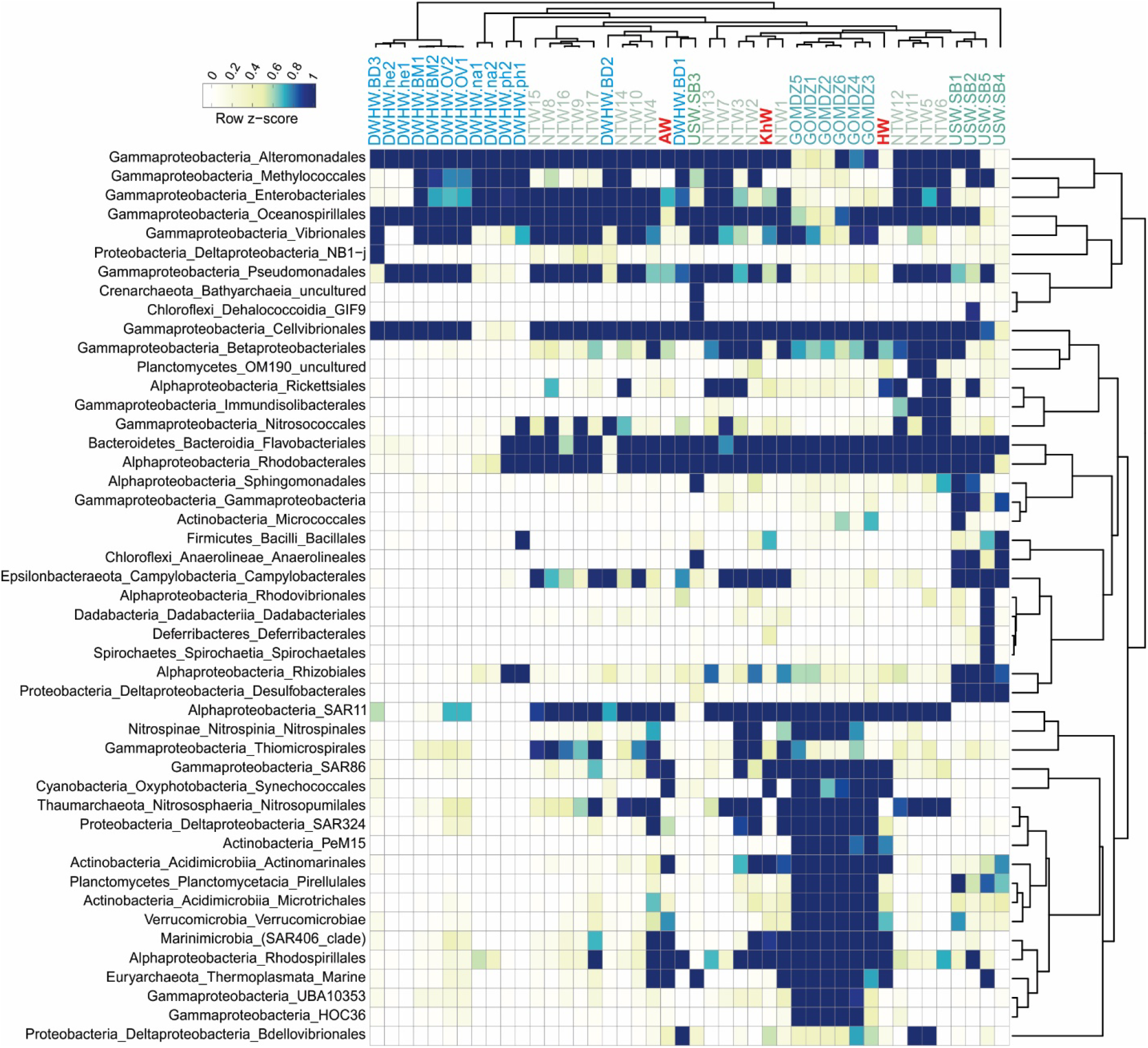
Prokaryotic community composition of PG water samples along with 41 metagenomes originating from oil-polluted marine water samples derived from the relative abundance of 16S rRNA gene in unassembled reads. Row names are at the order level. For taxa with lower frequency, higher taxonomic level is shown (47 taxa in total). Columns are the name of water samples. Samples are clustered based on Pearson correlation and the color scale on the top left represents the raw Z-score.

*Flavobacteriales* and *Rhodobacterales* were present in relatively high abundance in almost all oil-polluted samples except for those with recent pollution. NTW5, NTW6, NTW11, NTW12 samples incubated with MC252 oil for 32-64 days represent similar prokaryotic composition dominating taxa that are reportedly involved in degrading recalcitrant compounds like PAHs in the middle-to-late stages of oil degradation process (*Alteromonadales, Cellvibrionales, Flavobacteriales,* and *Rhodobacterales*). Whereas at the earlier contamination stages samples represent a different community composition with a higher relative abundance of *Oceanospirillales* (e.g., NTW8, NTW9, NTW15, NTW16 and NTW17 sampled after 0-8 days incubation).

The non-metric multidimensional analysis of the prokaryotic community of 106 oil-polluted water and sediment samples together with the PG samples is represented in **Figure 2**. The water and sediment samples expectedly represent distinct community compositions. AW sample is placed near samples treated with phenanthrene and naphthalene in the NMDS plot showing the impact of aromatic compounds on its microbial community. KhW sample is located near NTW13 in the plot both of which have experienced recent oil pollution.

**Figure 2.**
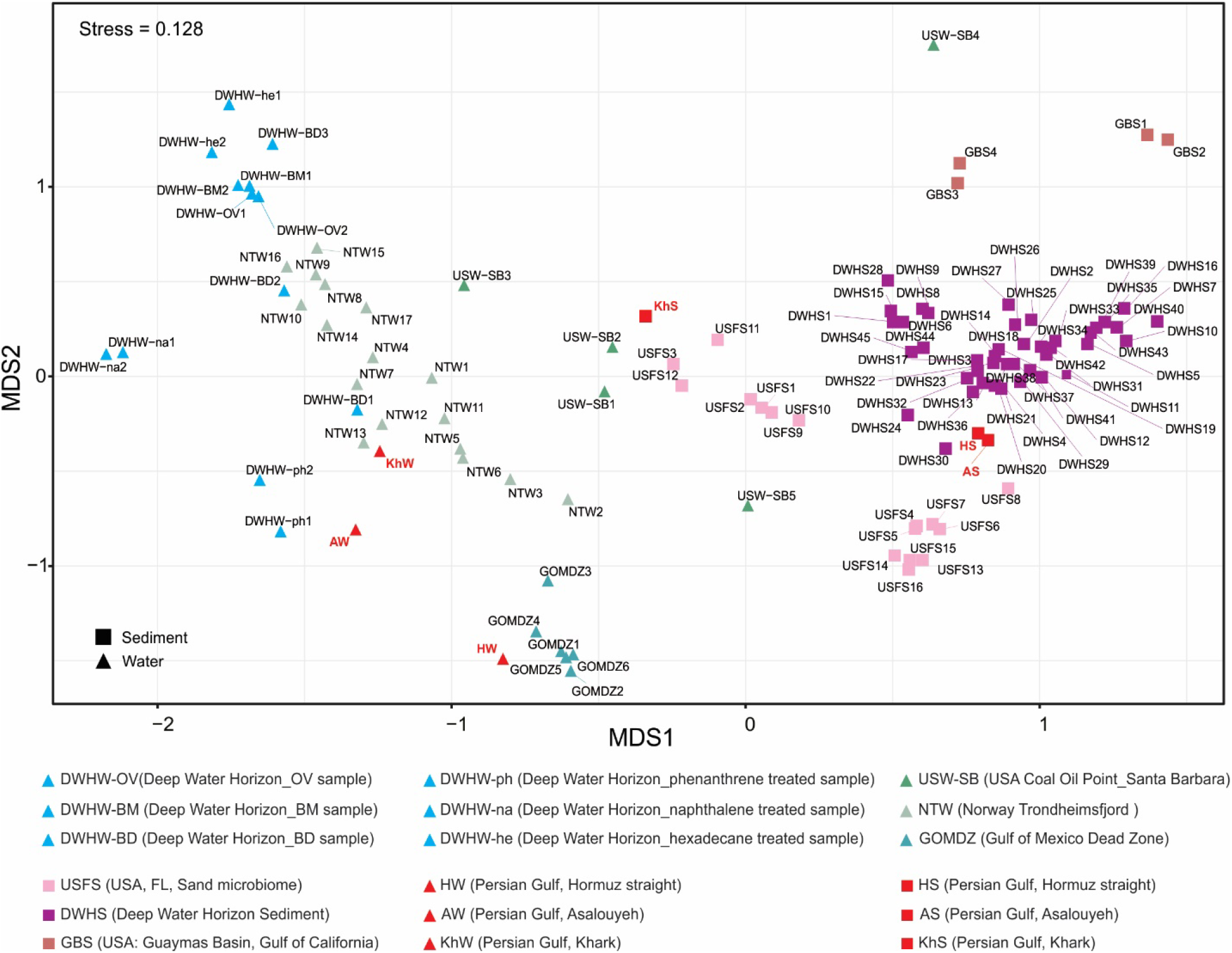
Non-metric multidimensional scaling (NMDS) of the Persian Gulf water and sediment metagenomes along with oil-polluted marine water and sediment metagenomes based on Bray-Curtis dissimilarity of the abundance of 16S rRNA gene in unassembled reads at the order level. Samples with different geographical locations are shown in different colors. PG water and sediment samples are shown in red. Water and sediment samples are displayed by triangle and square shapes respectively.

The orders *Oceanospirillales, Alteromonadales* and *Pseudomonadales* are present in relatively high abundances in all oil pollutes water samples except for HW (PG input water) and samples collected from the northern Gulf of Mexico dead zone (GOMDZ) (**Figure 1**). Persian Gulf is located in the proximity of the developing oxygen minimum zone (OMZ) of the Arabian Seas that is slowly expanding towards the Gulf of Oman (Bandekar *et al.*, 2018; Bertagnolli and Stewart, 2018). Potential water exchange with OMZ areas could be the cause of higher similarity to the GOMDZ microbial community (Thrash *et al.*, 2016).

While marine prokaryotes represent vertical stratification with discrete community composition along depth profile, the prokaryotic communities of the oil-polluted areas according to our analyses; are consistently dominated by similar taxa regardless of sampling depth or geographical location. We speculate that the high nutrient input due to crude oil intrusion into the water presumably disturbs this stratification and HC degrading microorganisms are recruited to the polluted sites where their populations flourish. The inherent heterogeneity of the sediment prokaryotic communities is retained even after exposure to oil pollution reflected in their higher alpha diversity (**Supplementary Figure S6**), however similar taxa dominate the community in response to oil pollution (**Figure 3**).

**Figure 3.**
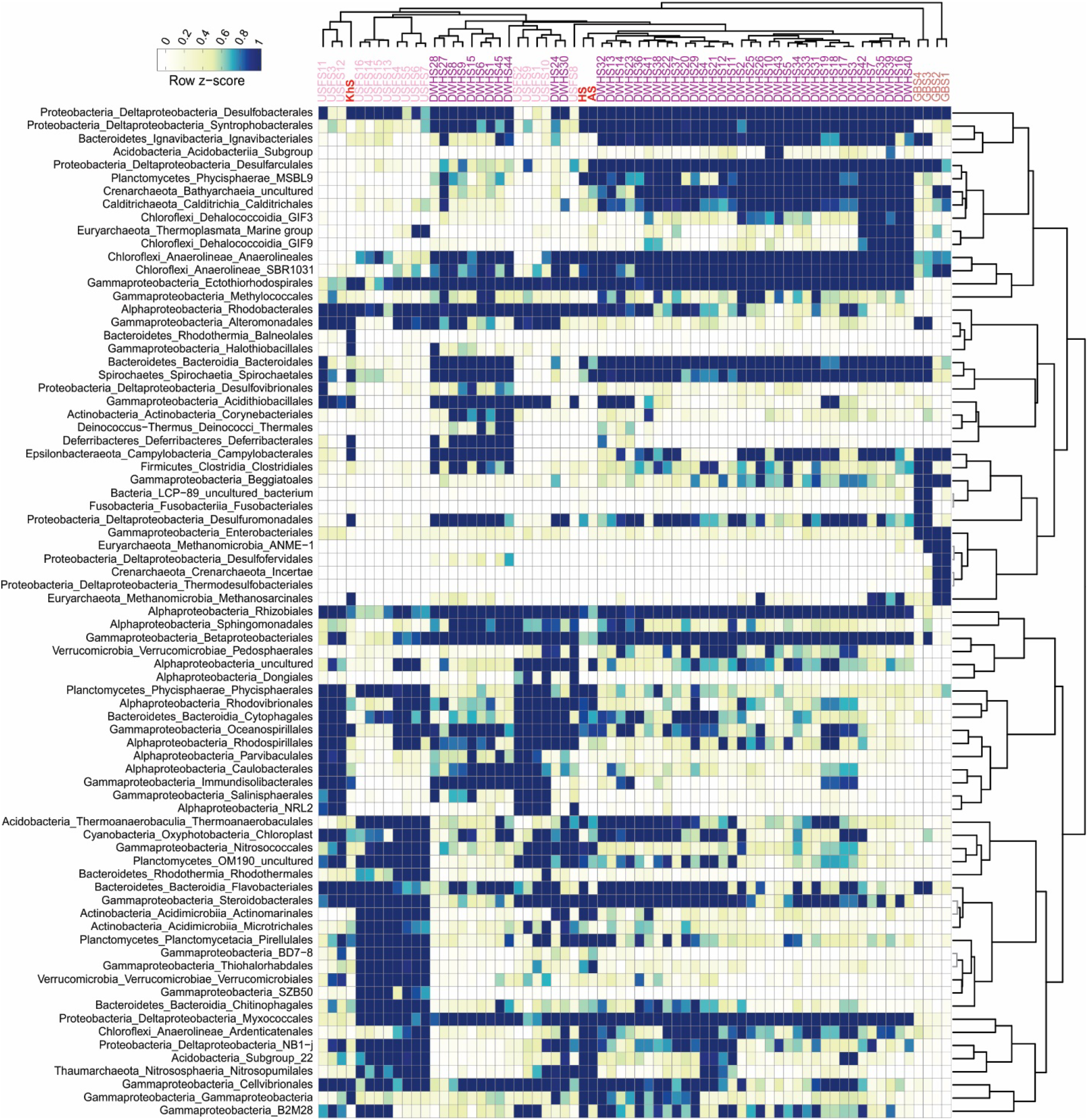
Prokaryotic community composition of PG sediment samples along with 65 metagenomes originating from oil-polluted marine sediment samples derived from the relative abundance of 16S rRNA gene in unassembled reads. Row names are at the order level. For taxa with lower frequency, the higher taxonomic level is shown (77 taxa in total). Columns are the name of sediment samples. Samples are clustered based on Pearson correlation and the color scale on the top left represents the raw Z-score.

In sediment samples, *Deltaproteobacteria* had the highest abundance followed by *Gammaproteobacteria* representatives. *Ectothiorhodospirales, Rhizobiales, Desulfobacterales, Myxococcales* and *Betaproteobacteriales* representatives were present in almost all samples at relatively high abundances (**Figure 3**). Sulfate/nitrate-reducing bacteria are major HC degraders in sediment showing substrate specificity for anaerobic HC degradation (Abbasian *et al.*, 2015, 2016). *Desulfobacterales* and *Myxococcales* are ubiquitous sulfate-reducers, present in almost all oil-polluted sediment samples (Paissé *et al.*, 2008; Stagars *et al.*, 2017). Sulfate-reducing *Deltaproteobacteria* play a key role in anaerobic PAH degradation especially in sediments containing recalcitrant HC types (Davidova *et al.*, 2018). Members of *Rhizobiales* are involved in nitrogen fixation which accelerates the HC removal process in the sediment samples (Shin *et al.*, 2019), therefore their abundance increase in response to oil pollution (**Figure 3**).

Prokaryotes involved in nitrogen/sulfur cycling of sediments are defined by factors such as trace element composition, temperature, pressure and more importantly, depth and oxygen availability. In oil-polluted sediment samples, the simultaneous reduction of available oxygen with an accumulation of recalcitrant HCs along the depth profile complicates the organic matter removal. However, anaerobic sulfate-reducing HC degrading bacteria will cope with this complexity (Acosta-González *et al.*, 2013). Our results show that sampling depth (surface or subsurface) of the polluted sediments defines the dominant microbial populations. The prokaryotic community of HS and AS samples represent similar phylogenetic diversity (**Figures 2 and 3**). Their prokaryotic community involved in the nitrogen and sulfur cycling resembles the community of DWHS samples. KhS sample has a similar prokaryotic community to deeper sediment samples collected from 30-40cm depth (USFS3, USFS11, and USFS12) which could be due to our sampling method using a grab device. The co-presence of the orders *Methanosarcinals, Alteromanadales* and *Thermotogae* (*Petrotogales*) in KhS sample hint at potential oil reservoir seepage around the sampling site in the Khark Island since these taxa are expected to be present in oil reservoirs (Liu *et al.*, 2018). Hydrocarbon degrading microbes show ubiquitous distribution in almost all oil-polluted water and sediment samples and their community varies depending on the type of oil pollution present at the sampling location and the exposure time. It appears that the sediment depth and oil contaminant type, define the microbial community of sediments.

### Genome-resolved metabolic analysis of the Persian Gulf’s prokaryotic community along the pollution continuum

A total of 82 metagenome-assembled genomes (MAGs) were reconstructed from six sequenced metagenomes of the PG (completeness ≥40% and contamination ≤5%). Amongst them, eight MAGs belonged to domain Archaea and 74 to domain bacteria. According to GTDB-tk assigned taxonomy (release89), reconstructed MAGs were affiliated to *Gammaproteobacteria* (36.6%), *Alphaproteobacteria* (12.2%), *Flavobacteriaceae* (9.7%), *Thermoplasmatota* (5%) together with some representatives of other phyla (MAG stats in **Supplementary Table S4**).

A collection of reported enzymes involved in the degradation of different aromatic and aliphatic hydrocarbons under both aerobic and anaerobic conditions were surveyed in the annotated MAGs of this study (Pérez-Pantoja *et al.*, 2010; Abbasian *et al.*, 2015, 2016; Meckenstock *et al.*, 2016; Rabus *et al.*, 2016; Espínola *et al.*, 2018). KEGG orthologous accession numbers (KOs) of genes involved in HC degradation were collected and the distribution of KEGG orthologues detected at least in one MAG (n=76 genes) is represented in **Figure 4**.

**Figure 4.**
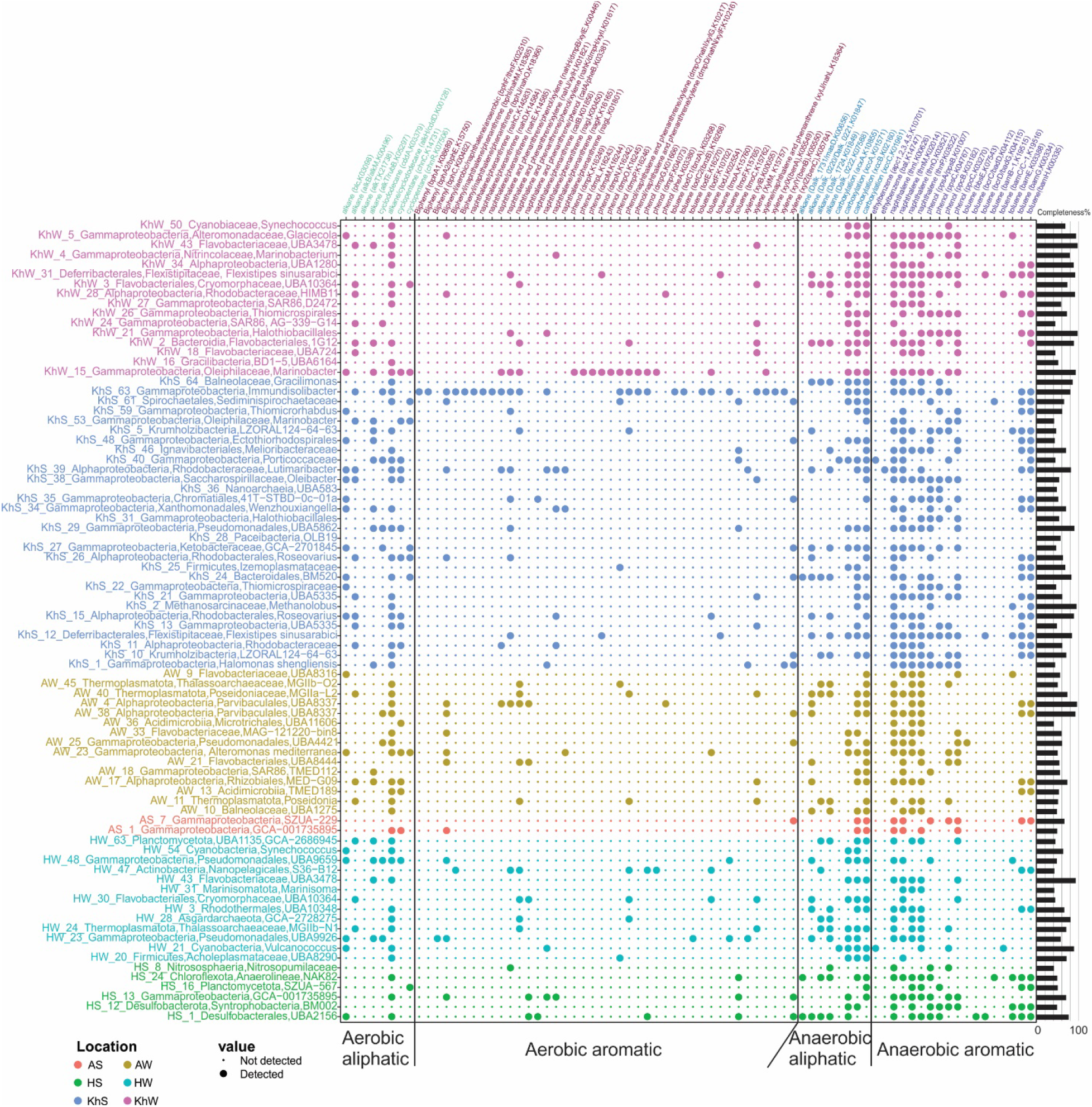
Hydrocarbon degrading enzymes present in recovered MAGs from the PG water and sediment metagenomes with completeness higher than 40% and contamination lower than 5%. Row names represent the taxonomy of recovered MAGs and their completeness is provided as a bar plot on the right side. The color indicates the MAG origin. The size of dots indicates the presence or absence of each enzyme in each recovered MAG. Columns indicate the type of hydrocarbon and in the parenthesis is the name of the enzyme hydrolyzing this compound followed by its corresponding KEGG orthologous accession number.

A combination of different enzymes runs the oil degradation process. Mono- or dioxygenase enzymes, are the main enzymes triggering the degradation process under aerobic conditions. Under anaerobic conditions, degradation is mainly triggered by the addition of fumarate or in some cases by carboxylation of the substrate. Therefore, bacteria containing these genes will potentially initiate the degradation process that will be continued by other heterotrophs. Enzymes such as decarboxylase, hydroxylase, dehydrogenase, hydratase and isomerases through a series of oxidation/reduction reactions act on the products of initiating enzymes mentioned above.

Various microorganisms with different enzymatic capability cooperate for cleavage of hydrocarbons into simpler compounds that could enter common metabolic pathways. Mono-dioxygenases which are involved in the degradation of alkane (alkane 1-monooxygenase, alkB/alkM), cyclododecane (cyclododecanone monooxygenase, cddA), Biphenyl (Biphenyl 2, 3-dioxygenase subunit alpha/beta, bphA1/A2, Biphenyl-2, 3-diol 1, 2-dioxygenase, bphC), phenol (phenol 2-monooxygenase, pheA), toluene (benzene 1, 2-dioxygenase subunit alpha/beta todC1/C2, hydroxylase component of toluene-4-monooxygenase, todE), xylene (toluate/benzoate 1,2-dioxygenase subunit alpha/beta/electron transport component, xylX/Y/Z, hydroxylase component of xylene monooxygenase, xylM) and naphthalene/phenanthrene (catechol 1,2 dioxygenase, catA, a shared enzyme between naphthalene/phenanthrene /phenol degradation) were detected in recovered MAGs of the PG.

The key enzymes including Alkylsuccinate synthase (I)/(II) (assA1/A2), benzylsuccinate synthase (BssA)/benzoyl-CoA reductase (BcrA), ethylbenzene dehydrogenase (EbdA) and 6-oxo-cyclohex-1-ene-carbonyl-CoA hydrolase (BamA) that are responsible for the degradation of alkane, toluene, ethylbenzene and benzoate exclusively under anaerobic conditions were not detected in reconstructed MAGs of this study. This shows that recovered MAGs of this study are not initiating anaerobic degradation via known pathways while they have the necessary genes to continue the degradation process started by other microorganisms.

Exploring the distribution of aromatic/aliphatic HC degrading enzymes within our MAGs highlights *Alphaproteobacteria* and *Gammaproteobacteria* representatives as the most prominent degraders in PG water and sediment (Figure 4). MAG KhS_63 affiliated to *Immundisolibacter* contains various types of mono-dioxygenases and is capable of degrading a diverse range of hydrocarbons such as alkane, cyclododecane, toluene and xylene. Members of this genus have been shown to degrade high molecular weight PAHs (Corteselli *et al.*, 2017).

*Lutimaribacter* representatives have been isolated from seawater and reported to be capable of degrading cyclohexylacetate (Iwaki *et al.*, 2013). We also detect enzymes responsible for alkane, cycloalkane (even monooxygenase enzymes), and naphthalene degradation under aerobic conditions as well as alkane, ethylbenzene, toluene and naphthalene degradation under anaerobic conditions in KhS_39 affiliated to this genus (**Figure 4**).

MAGs KhS_15 and KhS_26 affiliated to *Roseovarius* have the enzymes for degrading alkane (alkane monooxygenase, aldehyde dehydrogenase), cycloalkane, naphthalene and phenanthrene under aerobic and toluene and naphthalene under anaerobic condition. PAHs degradation has been reported for other representatives of this taxa as well (Shao *et al.*, 2015).

MAGs KhS_11 (a representative of *Rhodobacteraceae*) and KhS_53 (*Marinobacter*) have alkB/alkM, KhS_27 (GCA-2701845), KhS_29 (UBA5862) and KhS_40 (from *Porticoccaceae* family) have cddA, KhS_13 and KhS_21 (UBA5335) and KhS_38 (*Oleibacter*) have both alkB/alkM and xylM genes and are among microbes that are initiating the degradation of alkane, cycloalkane and xylene compounds. Other MAGs recovered from Khark sediment are involved in the continuation of the degradation pathway. For example, KhS_1 is affiliated to the genus *Halomonas* and has different enzymes to degrade intermediate compounds. *Halomonas* representatives have been frequently isolated from oil-polluted environments (Barbato *et al.*, 2019). The phylum *Krumholzibacteria* has been first introduced in 2019 and reported to contain heterotrophic nitrite reducers (Youssef *et al.*, 2019). MAGs KhS_5 and KhS_10 are affiliated to this phylum and contain enzymes involved in anaerobic degradation of toluene, phenol, and naphthalene (**Figure 4**).

MAGs KhS_12 and KhW_31 affiliated to the genus *Flexistipes*, in *Deferribacterales* order, have been reconstructed from both KhW and KhS samples. *Deferribacterales* are reported to be present in the medium to high-temperature oil reservoirs with HC degradation activity and also in high-temperature oil-degrading consortia (Head *et al.*, 2014; Zhou *et al.*, 2019). The type strain of this species was isolated from environments with the minimum salinity of 3% and temperature of 45-50 °C (Fiala *et al.*, 1990). The presence of this genus in KhS could be due to natural oil seepage from the seabed as PG reservoirs mainly have medium to high temperature and high salinity. Representatives of *Deferribacterales* have been reported to be sulfate/sulfur-reducing HC degrading bacteria under anaerobic conditions (Gieg *et al.*, 2010). MAGs KhS_12 and KhW_31 do not encode genes for sulfate/sulfur reduction; however, they contain enzymes involved in the degradation of alkane, phenol, toluene and naphthalene under anaerobic conditions.

As mentioned earlier, *Flavobacteriales* are potent marine indigenous hydrocarbon degraders that bloom in response to oil pollution, especially in water samples (Liu and Liu, 2013). *Flavobacteriales* affiliated MAGs (KhW_2, KhW_3, AW_21, and AW_33) were recovered from polluted water samples of KhW and AW and mostly contain enzymes that participate in the degradation of aromatic compounds under anaerobic conditions. KhW_2 and KhW_3 also have both alkB/M (alkane monooxygenase) and xylM enzyme, which initiates the bioremediation process of alkane and xylene in Khark water. Among other recovered MAGs from KhW sample, KhW_18 (UBA724), KhW_24 (clade SAR86), KhW_43 (UBA3478) have alkB/M and xylM, KhW_24 (clade SAR86) have alkB/M and cddA, and KhW_28 (from *Rhodobacteraceae* family) have alkB/M and pheA genes in their genome to initiate the degradation process (**Figure 4**).

*Marinobacter* (KhW_15) is another potent MAG reconstructed from KhW sample. It has been frequently reported that this genus is one of the main cultivable genera that play a key role in bioremediation of a wide range of oil derivatives, especially in marine polluted ecosystems (Gutierrez *et al.*, 2013; Barbato *et al.*, 2019).

Marine Group II (MGII) and *Poseidonia* representatives of *Thermoplasmatota* that have been reported to be nitrate-reducing *Archaea* (Rinke *et al.*, 2019), were recovered from AW sample (AW_40, AW_45) and contain several enzymes contributing in alkane (alkane monooxygenase, aldehyde dehydrogenase) and naphthalene/phenanthrene/phenol/xylene degradation (decarboxylase) under aerobic conditions. The HC degradation potential of representatives of these taxa has been previously reported (Redmond and Valentine, 2012; Jeanbille *et al.*, 2016).

In Asalouyeh water sample, MAGs AW_25 (UBA4421) and AW_38 (UBA8337) have cddA, AW_21 (UBA8444) have catA, AW_11 (*Poseidonia*) and AW_17 (from *Rhizobiales* order) have both alkB/M and xylM, and AW_4 (UBA8337) have catA and pheA genes and trigger the breakdown of their corresponding oil derivatives. Other recovered genomes will act on the product of initiating enzymes. For instance, AW_23 contains enzymes involved in the degradation of naphthalene, phenol and cyclododecane and is affiliated to genus Alteromonas (**Figure 4**).

Three recovered MAGs of HW affiliated to *Pseudomonadales* (HW_23), *Poseidoniales* (HW_24) and *Flavobacteriales* (HW_30) contain some initiating enzymes to degrade cyclododecane/biphenyl/toluene, alkane/xylene and alkane/xylene/naphthalene/phenanthrene respectively. A representative of *Heimdallarchaeia* that are mainly recovered from sediment samples was reconstructed from the Hormuz water sample. The HW_28 MAG has completeness of 81% and contains enzymes involved in anaerobic degradation of alkanes. This archaeon could potentially be an input from the neighboring OMZ as this phyla contain representatives adopted to microoxic niches (Bulzu *et al.*, 2018). Having genes with the potential to initiate the oil derivative degradation in the input water with no oil exposure, reiterates the intrinsic ability of marine microbiota for oil bioremediation.

## Conclusions

Exploring the response of the marine microbial communities to oil pollutions has received an increasing attention over the last decade specifically after the “Deepwater Horizon oil spill”. However, the influence of long-term exposure to oil derivatives in ecosystems such as Persian Gulf that hosts almost half of the world’s oil reserves and has been chronically exposed to recurrent natural and accidental oil pollutions has remained entirely unknown. Understanding the microbial dynamics in response to oil pollution at different locations of the Persian Gulf can function as a valuable model system for advancing our knowledge and preparedness for managing oil spill accidents in the future.

Our extensive analysis of available oil-polluted water and sediment metagenomes (*n*=106) together with the Persian Gulf samples (*n*=6) show that the chronic exposure to trace amounts of oil derivatives has altered the microbial community of the Persian Gulf. Even though the pollution remained below our detection limit of 50 μg/l, the long-standing trace oil pollution imposes a consistent selection pressure on the microbial community of the input water selecting for oil-degrading microbes capable of degrading major local pollutants.

Our results show that certain members of the marine community (e.g., representatives of *Oceanospirillales, Flavobacteriales, Alteromonadales, Rhodobacterales Cellvibrionales* along with nitrate/sulfate-reducing HC degrading microbes residing in sediments) are the main drivers of oil degradation in water and sediment samples and regardless of the water column depth and initial composition, the microbial community converges in response to the pollution. It seems that the microbes capable of degrading more labile components of the pollutant will be recruited to the pollution zone and their population will experience a bloom which will be followed by the next populations capable of degrading more recalcitrant components. These microbes employ an intricate division of labor in initiating and carrying out different stages of the bioremediation process. Higher-resolution spatiotemporal analysis of the microbial community of this highly heterogeneous ecosystem in future studies can reveal important ecological adaptations to oil pollutants.

## Experimental Procedures

### Sampling site description, sample collection and DNA extraction

Three different locations of the Persian Gulf (abbreviated as PG) were selected for sampling based on their different level of potential exposure to oil pollution. PG receives its major marine water input from the Indian Ocean through the Strait of Hormuz in spring (peaking around May-June) while the Arvandrood river delta in the northwest feeds the Gulf with freshwater input (Al Azhar *et al.*, 2016). Sampling locations were selected based on their potential level of exposure to the oil derivatives contamination and sampled in spring and summer 2018. Water body around Hormuz Island (27.06112 N, 56.24636 E) is the representative of input water whereas water bodies around Asaluyeh county in Bushehr province (27.31767 N, 52.32542 E) and Khark Island (29.13194 N, 50.18105 E) are exposed to oil derivatives contaminants due to their position as a petrochemical and industrial hub or oil export center (Akhbarizadeh *et al.*, 2016; Delshab *et al.*, 2016).

Water and sediment samples were collected using Niskin bottle and grab, respectively. Salinity, pH, dissolved oxygen content, conductivity and temperature of each sample were measured by HQ40D Portable Multi Meter (HACH).

Total petroleum hydrocarbon (TPH) content of water and sediment samples was measured by GC-FID method (Adeniji *et al.*, 2017). PAH compounds of water and sediment samples were determined by GC-Mass and ISO 13877 techniques respectively (Poster *et al.*, 2006). Since the TPH amount of the Khark sediment sample was higher than 1μg/g, the carbon distribution of this sample was also determined by Simulated Distillation (GC-SimDis) method based on ASTM D2887 standard (Boczkaj *et al.*, 2011). Other elements and anions of each sample were also detected by ICP-Mass and common measurement methods. For each sampling point, 20 liters of water samples were collected from 5 meters depth, pre-filtered through 20 μm (Albet DP5891150, Germany) and 5 μm pore-size (Albet DP5895150, Germany) filters. Biomass was finally concentrated on 0.22 μm pore-size cellulose acetate filters (Sartorius 11107-142-N, Germany) using a peristaltic pump. Sediment samples were collected by grab and stored in separate falcon tubes. Water filters and sediment samples were kept on dry ice for transfer to the lab.

DNA from the water samples was extracted by a standard phenol-chloroform protocol as described elsewhere (Martín-Cuadrado *et al.*, 2007). DNA extraction for sediment samples was carried out using DNeasy PowerMax Soil DNA Extraction Kit (QIAGEN 12988-10, Germany) according to the manufacturer’s instruction. Extracted DNA samples were sequenced using Illumina Novaseq 6000 platform (PE150) (Novogene, Hong Kong).

### Estimation of 16S rRNA gene abundance by qPCR

*Halobacterium volcanii* (IBRC-M 10248) and *Escherichia coli* (IBRC-M 11074) were selected for drawing the standard curve as representative of domain archaea and bacteria, respectively. Their genomic DNA was (Marmur, 1961) and the 16S rRNA gene was amplified using universal primers (21F and 1492R for archaea and 27F and 1492R for bacteria)(Jiang *et al.*, 2006). DNA concentration was measured using Nanodrop (Thermo Nanodrop One/One-C Micro Volume Spectrophotometers) and copy number of double-strand DNA was estimated according to the formula: number of copies per μl = (concentration of PCR product (μl) * 6.022×10^23^) / (length of PCR product (bp) * 1×10^9^ * 650) in which, 650 is the molecular weight of one base pair in double-strand DNA and 6.022×10^23^ is Avogadro number. Domain-specific 16S rRNA primers named 338F (ACTCCTACGGGAGGCAGCAG), 533R (TTACCGCGGCTGCTGGCAC)(Mori *et al.*, 2013) and Parch519F (CAGCCGCCGCGGTAA), ARC915R (GTGCTCCCCCGCCAATTCCT)(Naghoni *et al.*, 2017) were selected to detect Bacteria and Archaea respectively. The qPCR reactions were performed using Power SYBR Green PCR Master Mix (BIOFACT, South Korea) in the MIC real-time PCR system (BioMolecular Systems, Australia). The unknown 16S rRNA copy number of each sample was calculated according to the standard curves (R^2^ value was higher than 99.0% in both curves). The total content of prokaryotes in each sample was calculated by the sum of 16S rRNA gene copies of bacteria and archaea.

### Reference metagenome collection

For comparative analyses, publicly available metagenomes deposited in the sequence read archive (SRA) of the Genebank were screened for the available metagenomic datasets originating from oil-polluted marine water and sediments. A total of 41 marine water and 65 marine sediment metagenomics datasets originating from oil-polluted samples were collected. Detailed description of these oil-polluted metagenomes is summarized in **Supplementary Table S3**. Water samples originate from Norway (Trondheimsfjord, *n*=17), Deepwater Horizon (Gulf of Mexico, *n*=13), northern part of the Gulf of Mexico (dead zone, *n*=6) and Coal Oil Point of Santa Barbara (*n*=5). Sediment samples originate from DWH Sediment (Barataria Bay, *n*=45), Municipal Pensacola Beach (USA, *n*=16) and a hydrothermal vent in Guaymas Basin (Gulf of California, *n*=4).

### Ribosomal RNA classification

A subset of 5 million reads was separated from each dataset and the reads affiliated to ribosomal RNA genes (16S/18S) were assigned using SSU-ALIGN (Nawrocki, 2009). Putative prokaryotic 16S rRNA sequences were blasted against the SILVA reference database (release 132SSUParc) and their taxonomic affiliation was assigned based on their closest hit if the read was ≥ 90bp at the similarity threshold of ≥ 90.

Non-metric multidimensional scaling (NMDS) analysis of oil-polluted marine water and sediment metagenomic samples worldwide together with the PG samples was performed using vegan package in Rstudio based on Bray-Curtis dissimilarity of the abundance of unassembled 16S rRNA gene reads of metagenomes (order-level). Alpha diversity of samples was also measured by vegan package in Rstudio based on the Shannon-Wiener index.

### Sequence assembly, binning and annotation

Paired-end reads of each sequenced dataset were interleaved using reformat.sh and quality trimmed by bbduck.sh scripts of BBMap toolkit (Bushnell, 2014). All trimmed sequences of each dataset were assembled separately using MEGAHIT (k-mer list 49,69,89,109,129 and 149)(Li *et al.*, 2015). Only contigs ≥1kb were binned into metagenome assembly genomes (MAGs) based on their different mapping depth and tetranucleotide frequency using MetaBat2 software (Kang *et al.*, 2019). Contamination and completeness of each MAG were evaluated using CheckM and MAGs with completeness above 40% and contamination lower than 5% were considered for further analysis (Parks *et al.*, 2015). The taxonomic affiliation of bins was assigned using GTDB-tk (Parks *et al.*, 2018). Putative genes were predicted using Prodigal (Hyatt *et al.*, 2010) and preliminarily annotated using Prokka in the metagenomics mood (Seemann, 2014). Predicted protein sequences of each MAG were further annotated using eggNOG-mapper (Huerta-Cepas *et al.*, 2018) and PfamScanner (Finn *et al.*, 2016).

**Figure 5.**
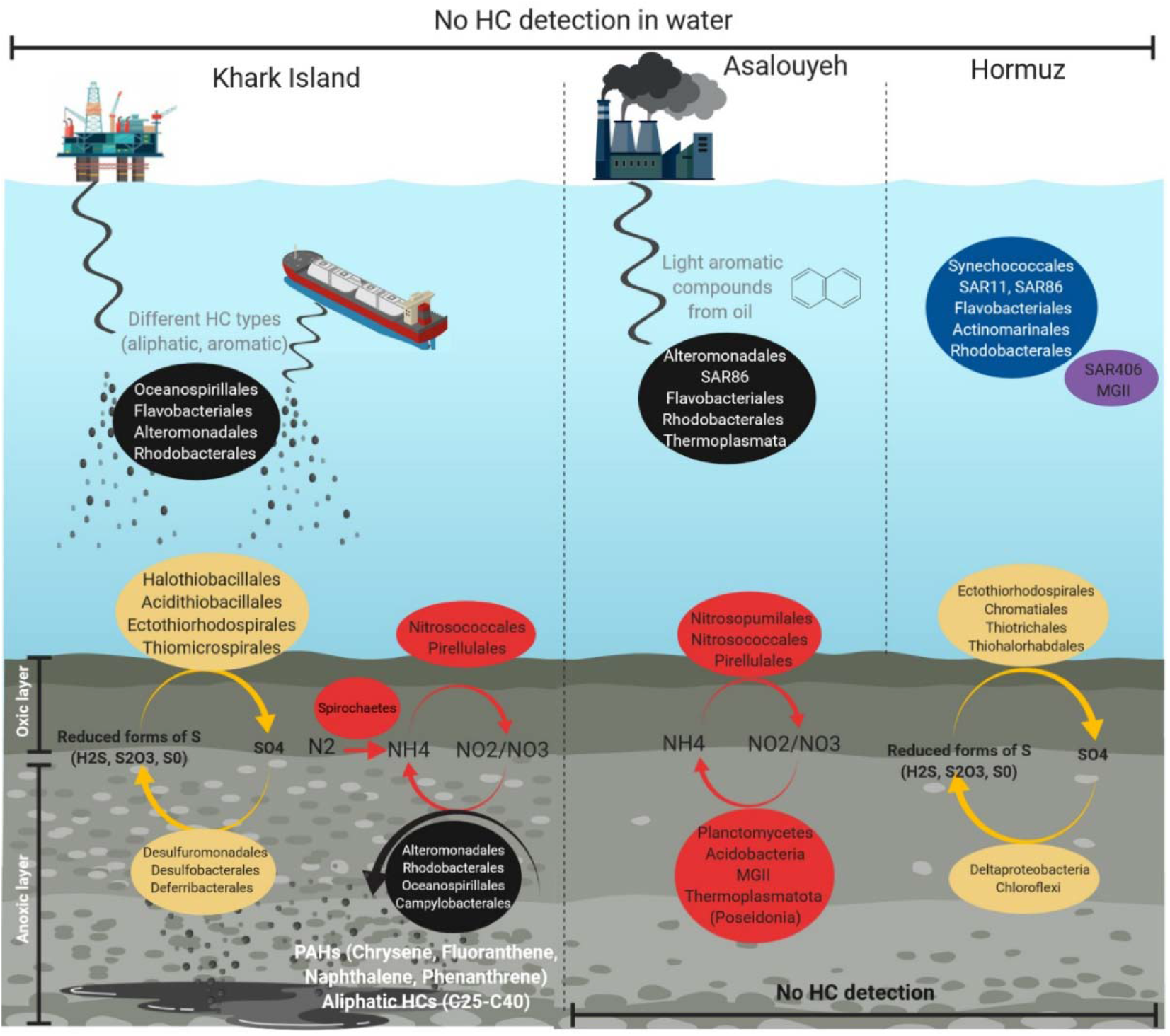
The microbial community dynamics of the Persian Gulf water and sediment samples in response to oil pollution. Hormuz Island was considered as a control location with the least impact from oil pollution. Taxa written in the blue frame are prevalent marine representatives present in HW. Microbial taxa in the Purple frame are mainly detected in OMZ areas and are also present in HW. Samples collected from Asalouyeh province are exposed to potential pollution caused by Gas field wastes. High oil trafficking, oil exploration and extraction, and natural oil seepage the main potential pollution sources in Khark Island. The potential pollutant types are shown in gray however the hydrocarbon pollution was below the detection limit in collected water samples. Black circles represent microorganisms that are involved in HC degradation in water samples from Asalouyeh and Khark Island. Microbes involved in sulfur and nitrogen cycle are shown in yellow and Red respectively. HS and AS have similar silt and sand-sized sediments with HC bellow the detection limit. KhS has gravel-sized particles and showed the highest oil pollution shown in white. “Figure created with BioRender.com”

## List of abbreviations

PG: Persian Gulf
HC: hydrocarbon
PAHs: Polyaromatic hydrocarbons

## Data availability

The metagenomic Raw read files of the Persian Gulf water and sediment samples as well as all the metagenome-assembled genomes (MAGs) reconstructed (Supplementary Table S5) in this study are archived at the DDBJ/EMBL/GenBank and can be accessed under the Bioproject PRJNA575141.

## Conflict of interests

The authors declare no competing interests.

## Authors’ contributions

MAA, SMMD, MM, and MSh devised the study. MRS and SMMD collected and processed the samples. MRS, MM, and LGM performed the bioinformatics analysis with assistance from KK. MRS and MM drafted the manuscript. All authors read and approved the manuscript.

## Acknowledgments

This study was supported by a grant of young pioneer biotechnologists from Iran Council for the development of Biotechnology to SMMD and the Iranian National Science Foundation (INSF) (to MAA). M.M. acknowledges financial support from the Science for Life Laboratory. The computational analysis was performed at the Center for High-Performance Computing, School of Mathematics, Statistics, and Computer Science, University of Tehran.

## Supplementary Figures

**Supplementary Figure S1.**
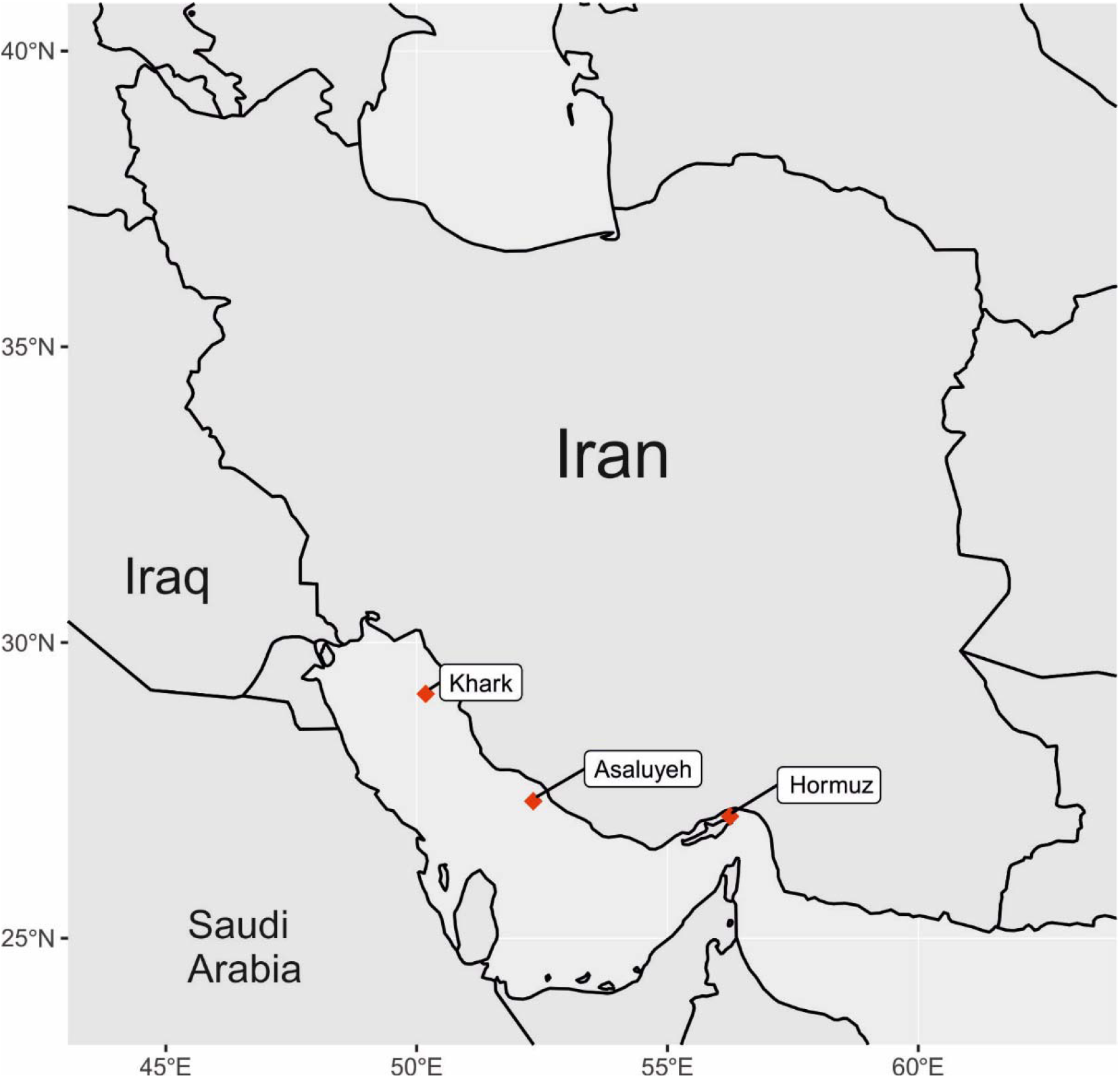
Sampling area and geographical location of three analyzed samples along the Persian Gulf pollution continuum. Hormuz Island as a probable unpolluted area located in the Persian Gulf water input, Asalouyeh province with mainly aromatic oil contaminants and Khark Island where mostly being polluted with different crude oil derivatives.

**Supplementary Figure S2.**
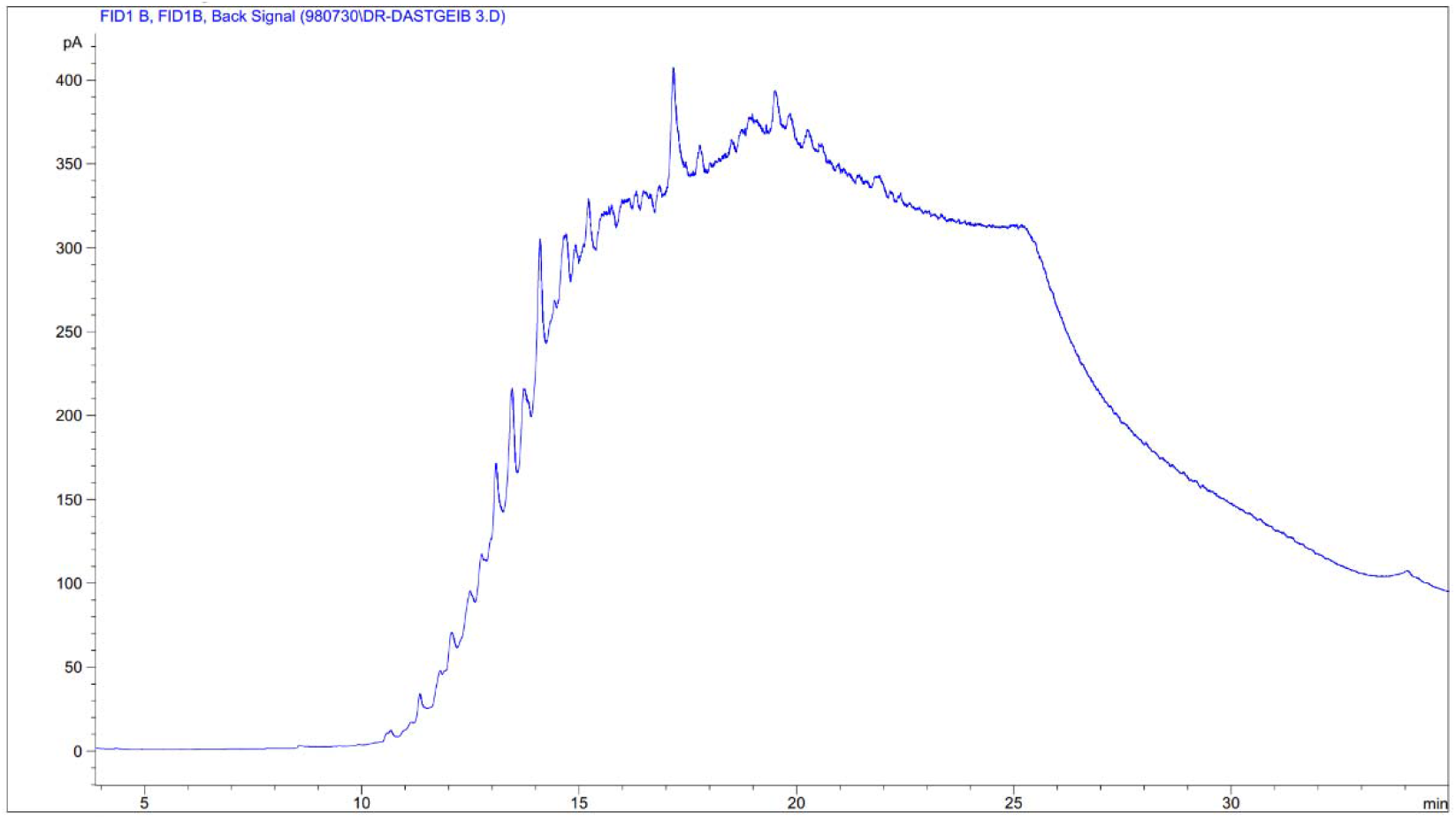
The carbon distribution of the bulk hydrocarbon compounds extracted from KhS sample measured by GC-SimDis method.

**Supplementary Figure S3.**
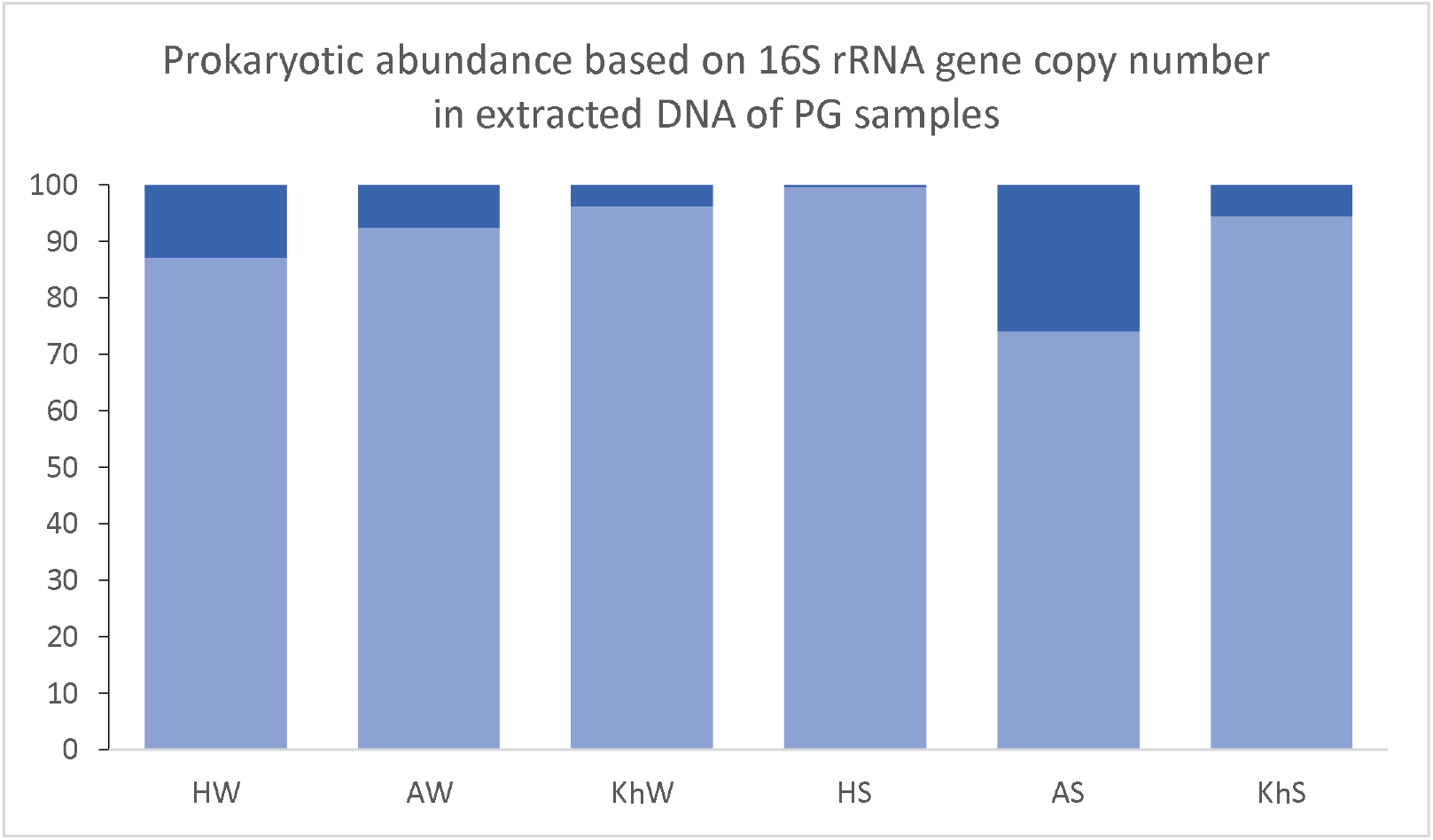
Prokaryotic 16S rRNA gene copy number abundance measured by qPCR presented as percentage.

**Supplementary Figure S4.**
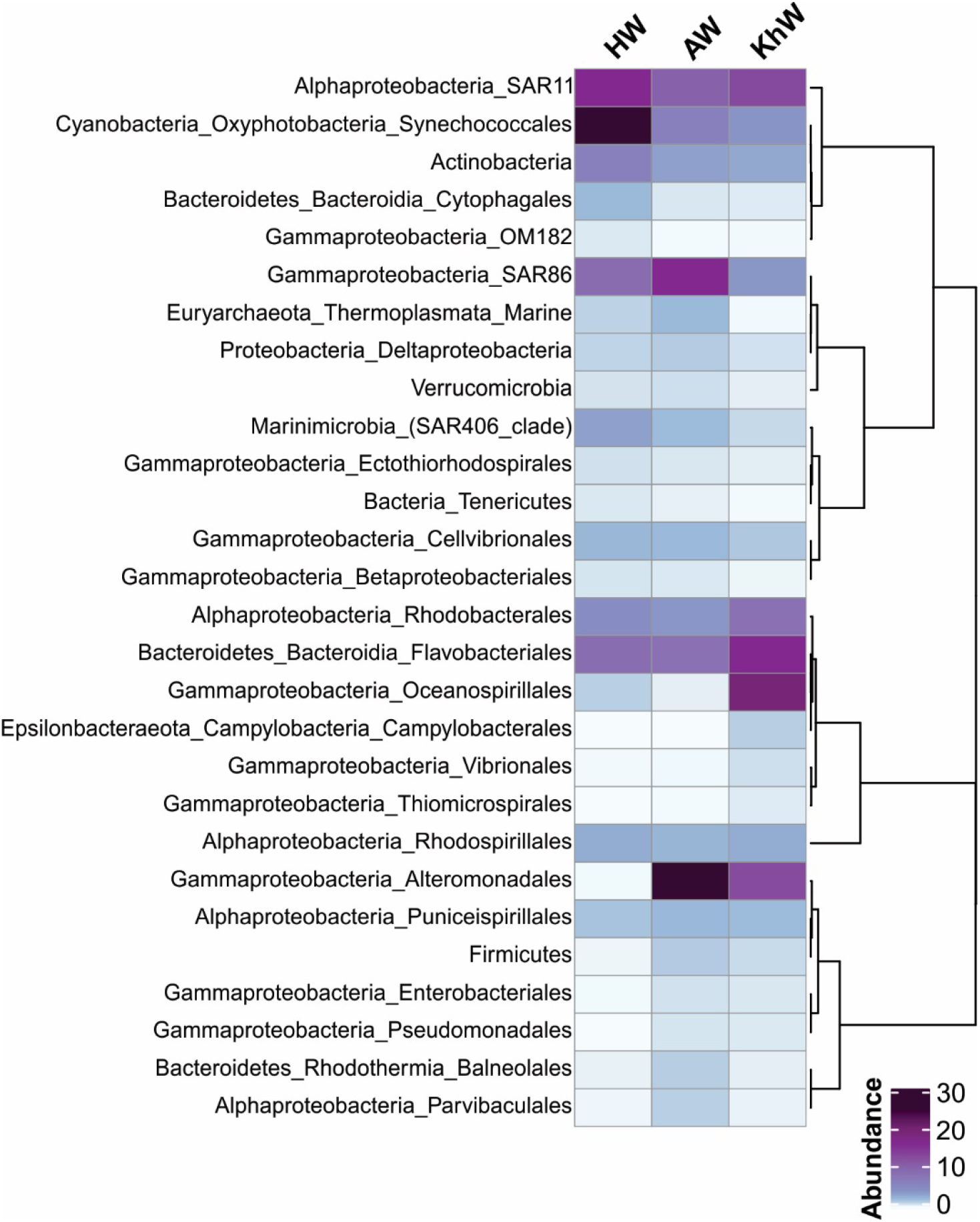
Prokaryotic community composition of the Persian Gulf water samples according to the abundance of 16S rRNA gene in unassembled reads. Row names are in order level. For some taxa with lower frequency, the sum of orders is displayed in their corresponding higher taxonomic level. There are total number of 28 taxa by which, samples are compared. Columns are the name of water samples.

**Supplementary Figure S5.**
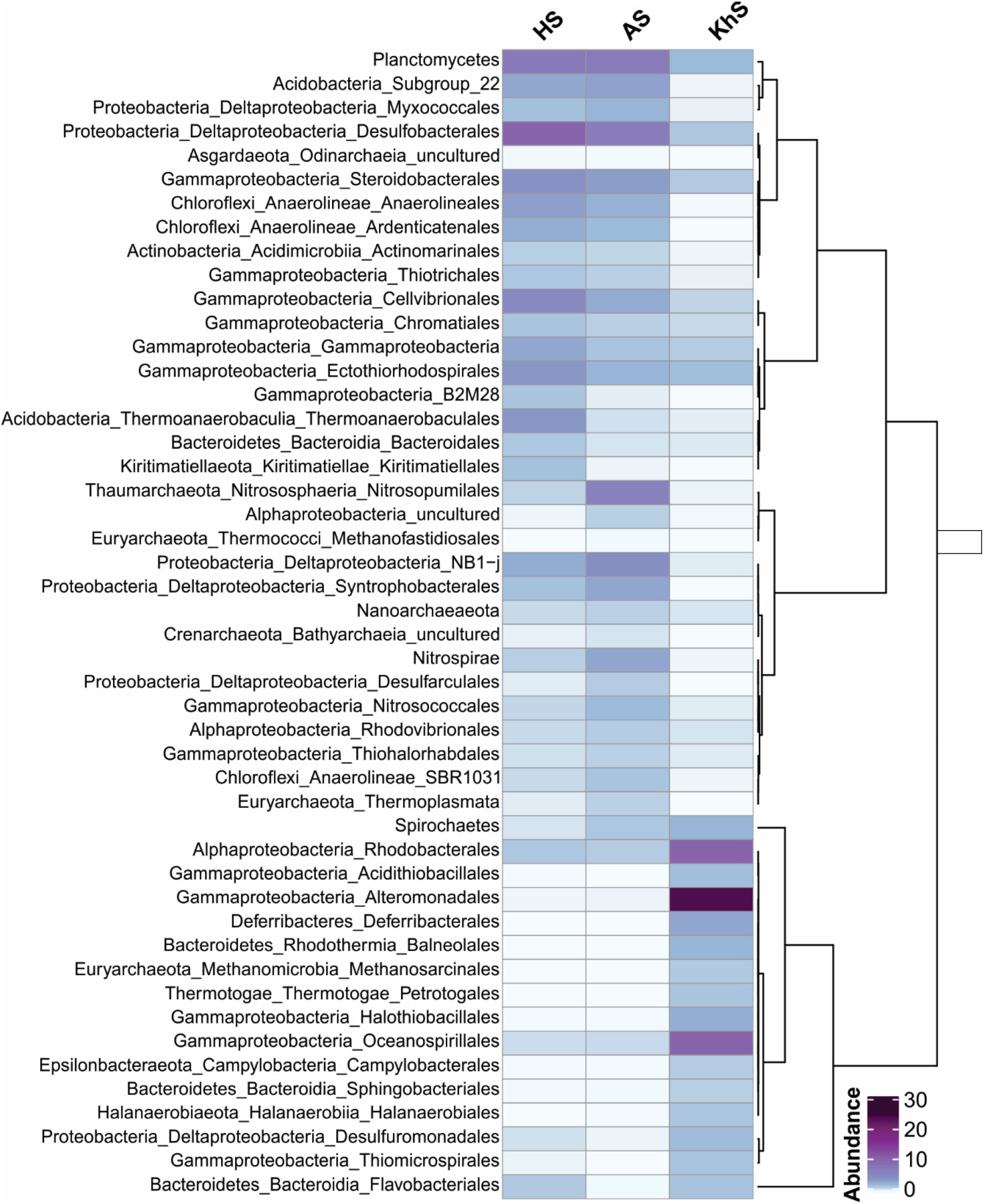
Prokaryotic community composition of the Persian Gulf sediment samples according to the abundance of 16S rRNA gene in unassembled reads. Row names are in the order level. For some taxa with lower frequency, the sum of orders is displayed in their corresponding higher taxonomic level. There are total number of 48 taxa by which, samples are compared. Columns are the name of sediment samples.

**Supplementary Figure S6.**
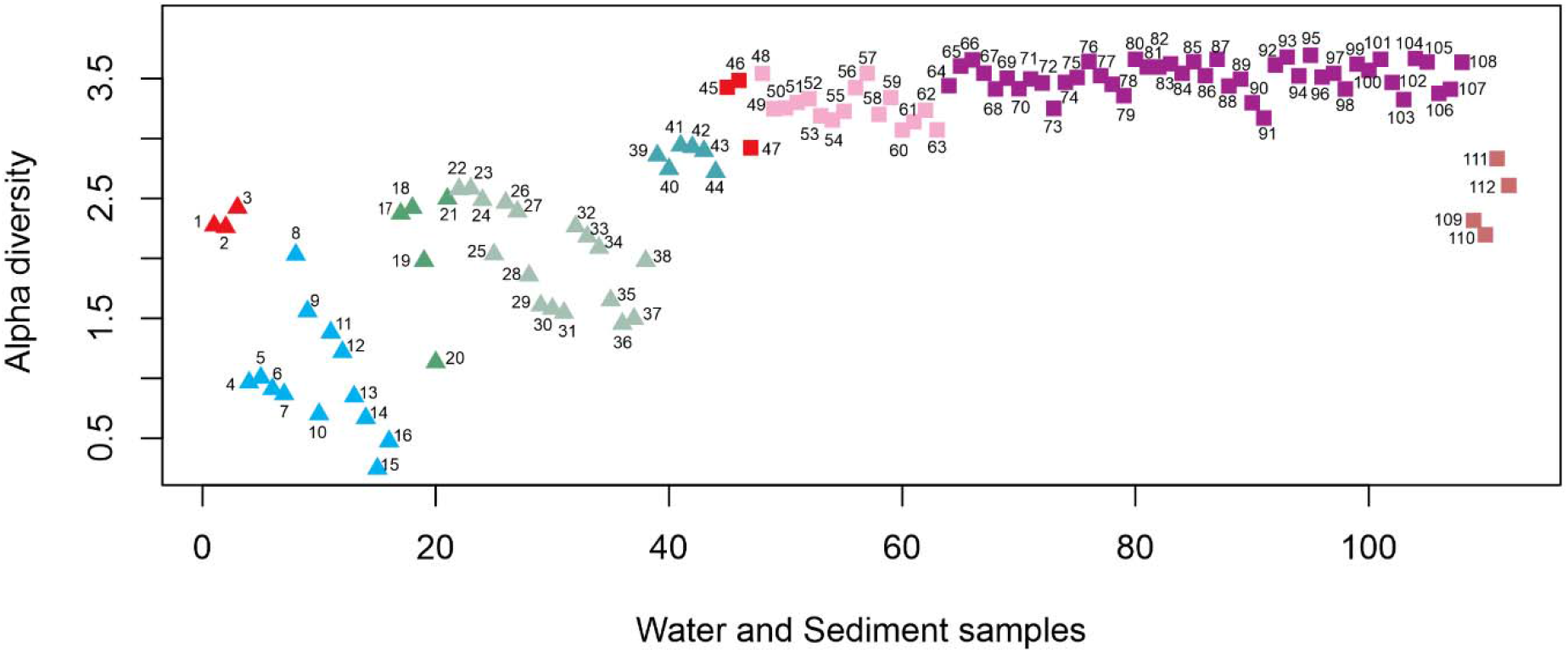
Alpha diversity of oil-polluted marine water and sediment sample together with the water and sediment samples collected from the Persian Gulf based on Shannon-Wiener index of the abundance of 16S rRNA gene in the unassembled reads clustered in th order level. Samples are color-coded as figure 1. Water and sediment samples are displayed by triangle and square shapes respectively.

## Notes

### Competing Interest Statement

The authors have declared no competing interest.

